# Analysing progress of SDG 6 in India: Past, Present, and Future

**DOI:** 10.1101/344184

**Authors:** Ajishnu Roy, Kousik Pramanick

## Abstract

Human endeavors to meet social and economic water needs at national scale might cause negative environmental manifestations and water stress from local to global scale. So, appropriation of Sustainable Development Goals requires a comprehensive monitoring and knowledge base of the water resource availability, utilization and access. Hence, scientific research progression has a significant role to facilitate the implementation of sustainable development goals through assessment and policy implementation from global to local scales. India holds a key position among developing economies with a complex interconnected web of a fast-growing population, coupled with biophysical stress, social deprivation and economic inequality related to water and sanitation. This study addresses some of these challenges related to monitoring and implementation of the targets of the United Nations Sustainable Development Goal 6 in India. Acknowledging the contribution of society and economy in sustainability paradigm, here we have chosen 28 indicators (clustered into eleven dimensions) under two major groups, concerning biophysical and social development aspects of water and sanitation. We have shown declining level of per capita biophysical water resource and slow to rapidly developing social indicators related to Sustainable Development Goal 6 in India. From past trends, we have calculated probable scenario of biophysical consumption of India up to 2050, which shows at least 1.3 times increase. This cumulative assessment framework contributes a tool to prioritize water resource appropriation, management response and policy implementations to national level sustainability of water and sanitation in India. We also advocate the necessity of restraining threats both at source and consumption process levels in order to ensure national water security for both human and biodiversity, keeping in mind the societal and economic development scenario.

## 1. Introduction

India is going through a time of deterioration of the environment, social deprivation and ineffective economy, combined with the rapidly growing population. This is an example of global scale initiation and progress of ‘Anthropocene’ (Steffen et al. 2007, 2011). During the 1980s, the concept of sustainable development was framed, which is “development that meets the needs of the present without compromising the ability of future generations to meet their own needs” (Brundtland Commission Report, 1987). In 1992 United Nations Conference on Environment and Development (UNCED, Rio de Janeiro Earth Summit), Agenda 21, calls for sustainable development indicators (SDIs) to “provide solid bases for decision-making at all levels and to contribute to a self-regulating sustainability of integrated environment and development system”. Emerging from Millennium Development Goals (MDGs), United Nations has set 17 Sustainable Development Goals (SDGs) and 169 targets in 2015. SDGs incorporate three pillars of sustainable development, i.e. environmental (climate action, life below water, life on land etc.), social (zero hunger, no poverty, gender equality, peace and justice and strong institutions etc.) and economic development (reduced inequalities, decent work and economic growth etc.) (Sachs, 2012).

There is a distinct trend that was set from the 1970s, especially from 1990s, that led to the contextual understanding and necessity for this study. It can be divided into two phases.

In the first phase, the significance of managing water resource sustainably was studied and advocated. Almost all of the works, concerned only with environmental aspect of water resource management. From the 1990s, the significance of the sustainable development of water resources has been increasing steadily. Biswas (1991, 1992) emphasised an analytical framework for drivers of the water crisis, socio-environmental considerations along with institutional responses for better management. Long before the present time, Serageldin (1995) had proposed ‘water resource management’ as a policy tool for sustainable future.

Gleick (1996) recommended that international organizations, national and local governments, with water providers adopt a ‘basic water requirement’ standard for human needs - 50 litres per person per day and promise access to this independently of an individual’s economic, social, or political status. In another work, Gleick (1998) proposed seven ‘sustainability criteria’ related to “basic water requirement (BWR)” to restructure long-term water planning. These criteria included – basic water requirement for the need and health of both human and ecosystems, water quality monitoring, institutional mechanism for water-related conflict resolution, water planning and decision making etc. Postel (2000) advocated a new water management paradigm which balances both ecological and economic functions of water by increasing water productivity (performing more functions with less water). Vörösmarty et al. (2000) have assessed the vulnerability of global water resource from both climate change and population growth. Jackson et al. (2001) analysed changes in water resources and projected future scenario for the USA. Vörösmarty et al. (2010) presented a global synthesis that jointly considers both human and biodiversity perspective on water security.

In the second phase, both economic and societal aspects started to get their due importance in important studies on sustainability analysis. In 2009, Rockström et al. devised a framework to understand if anthropogenic biophysical resource consumption has already exceeded or is going to surpass safe limits – ‘planetary boundaries’. This work marked the global onset of biophysical resource consumption monitoring in recent years. Then Steffen et al (2015) modified and improved the framework. One of the main problems in this work is that the indicators used in these works had not been corroborated completely with UN sustainable development goals. Also, the socioeconomic dimension was not comprehensively discussed. This was accomplished by Raworth (2012) through the incorporation of the social deprivation indicators. This acted as a complementary to planetary boundary framework and together formed ‘doughnut economy’ that touched all three pillars of sustainability – environment, economy and society. Water and sanitation are strongly related to public health too. Works in this area have also increased (Bartram et al, 2005, Moe and Rheingans, 2006, Montgomery and Elimelech, 2007, Bartram and Cairncross, 2010, Bartram et al. 2014). There has been an important discussion ongoing that specifically signifies water and sustainable development is perceived from this water-perspective (Bogardi et al. 2012, Madrid et al. 2013, Bhaduri et al. 2016, Ait-Kadi, 2016).

There has been some work on the appropriation of water resources and its management in India (Kumar et al. 2005, Shah and Koppen, 2006, Mall et al, 2006, Mujumdar, 2008). An important lacuna is that none of these incorporate socioeconomic dimensions in sustainability analysis of water resource. All of these studies logically imply the need for a national study on the sustainable development of water and sanitation that includes environment, society and economy together. That’s why this study was performed.

There is no national level study present focusing on both biophysical and socioeconomic aspects of SDG 6 i.e. sustainable development of water and sanitation. In this work, we have tried to find answers to the following: (1) How can we downscale SDG 6 indicators to a national scale more accurately? (2) How can we understand changes in dimensions of SDG 6 with time in order to contextualize their present values? (3) How can we use the past trends in per capita biophysical consumptions under SDG 6 to project probable future biophysical resource consumption at a national scale? This analysis measures the national performance of India on SDG 6 through11 dimensions, 28 indicators, segregated into 2 groups, biophysical and social development indicators, provides important findings of the relationship between biophysical resource use and well-being for India. Our work has been explained herein few steps. We first present our methodology, then results of our study on India; project probable future scenario of biophysical consumption for India; discuss limitations and scopes of this study, the applicability of SDG 6 as a tool in policymaking and necessities for future improvements in research.

## 2. Data and Method

We collected the data related to SDG 6 of India from Aquastat, FAOSTAT and World development indicators (world bank). As 3 main pillars of sustainable development are the environment, society, and economy, we also clubbed all SDG 6 associated indicators into 2 broad categories – biophysical and social development indicators (Fig. 1.), following planetary boundaries framework (Rockström et al, 2009; Steffen et al., 2015) and doughnut economics (Raworth, 2012, 2017).

**Fig. 1.**
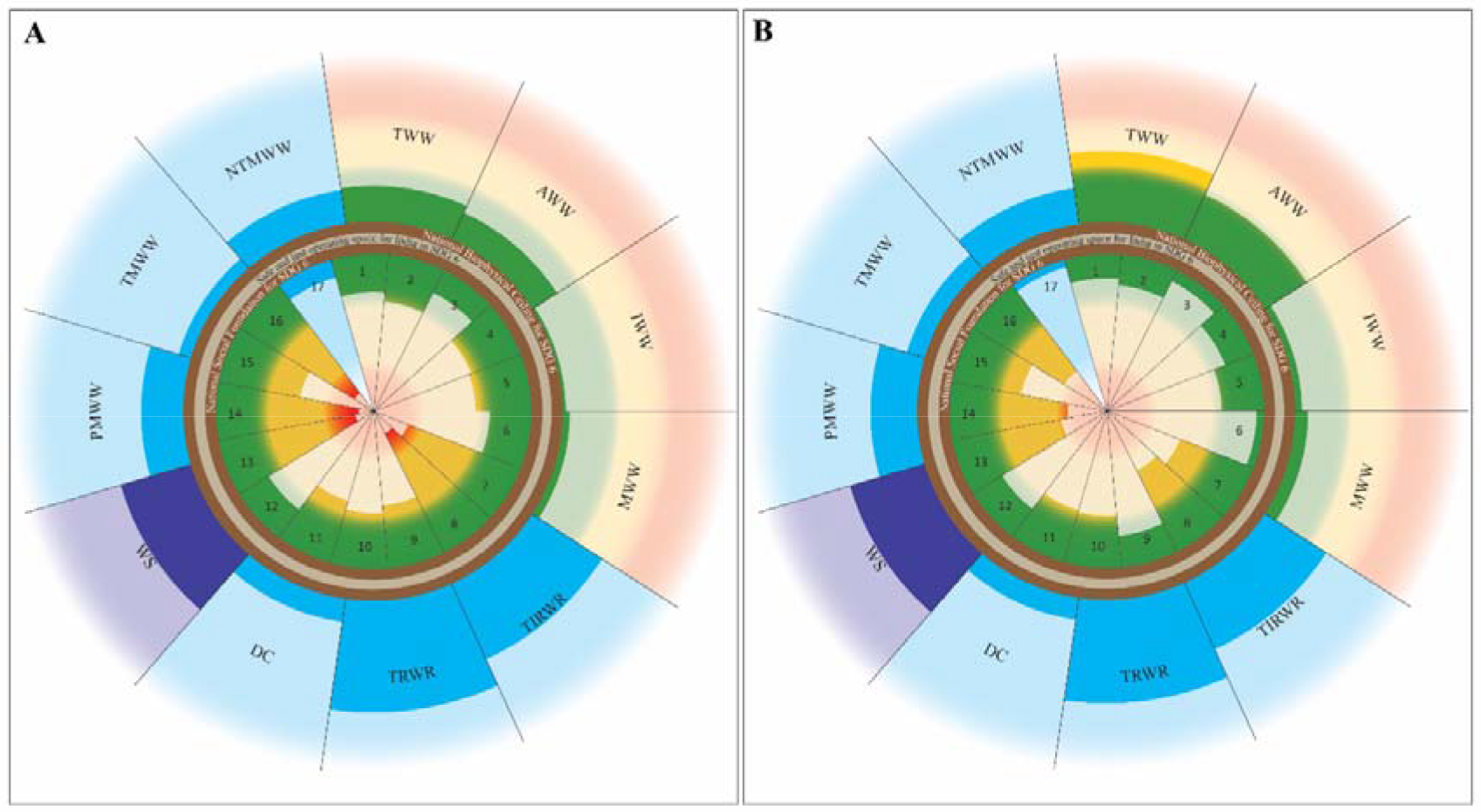
Changes in biophysical and social development indicators related to sustainable development goal 6 in India. A and B represent the status of sustainable development of India in 2000 and 2015 respectively. Eleven indicators of biophysical stress established over biophysical ceiling (outer chocolate ring) outwardly projected and seventeen indicators of social development established under social foundation (inner chocolate ring) inwardly projected for India. Biophysical indicators are – total water withdrawal (TWW), agricultural water withdrawal (AWW), industrial water withdrawal (IWW), municipal water withdrawal (MWW), total internal renewable water resource (TIRWR), total renewable water resource (TRWR), dam capacity (DC), water stress (WS), produced amount of municipal wastewater (PMWW), treated municipal wastewater (TMWW) and non-treated municipal wastewater (NTMWW). Indicators of social development are – (1) Improved sanitation facilities, (2) Improved sanitation facilities in rural areas, (3) Improved sanitation facilities in urban areas, (4) Improved water source, (5) Improved water source in rural areas, (6) Improved water source in urban areas, (7) People practicing open defecation, (8) Rural people practicing open defecation, (9) Urban people practicing open defecation, (10) People using at least basic drinking water services, (11) Rural people using at least basic drinking water services, (12) Urban people using at least basic drinking water services, (13) People using at least basic sanitation services, (14) Rural people using at least basic sanitation services, (15) Urban people using at least basic sanitation services, (16) Rural people using safely managed drinking water services, (17) Water use efficiency in agriculture. Green indicates safe operating space for biophysical indicators and thresholds for indicators of social development. Yellow indicates the zone of increasing impact for biophysical indicators and zone of increasing deprivation for indicators of social development. Red indicates the zone of high risk of serious impact for biophysical indicators and zone of high level of deprivation for indicators of social development. Light blue represents indicators without any boundary or thresholds (Unit – m^3^). Violet represents water stress (Unit - %). The area between the chocolate rings is the safe and just operating space for the sustainable development of water and sanitation in India.

### 2.1. Biophysical Indicators

We analyzed 11 indicators under 4 dimensions that indicate biophysical aspects of SDG 6 (Table 1). These 4 dimensions of biophysical indicators are - water withdrawal (4 indicators), water availability (3 indicators), wastewater (3 indicators) and water stress, WS (1 indicator). We calculated every biophysical indicator on per capita basis by dividing total values with population data of India, available from FAOSTAT database (except – water stress, expressed in percentage, %). We tried to include indicators for water pollution, water scarcity, the condition of water-related ecosystems, water footprint etc. Either no unanimously accepted indicator is available or there is no established database with annual data for the significantly longer duration. So, we could not include any indicator representing these dimensions.

**Table 1.**
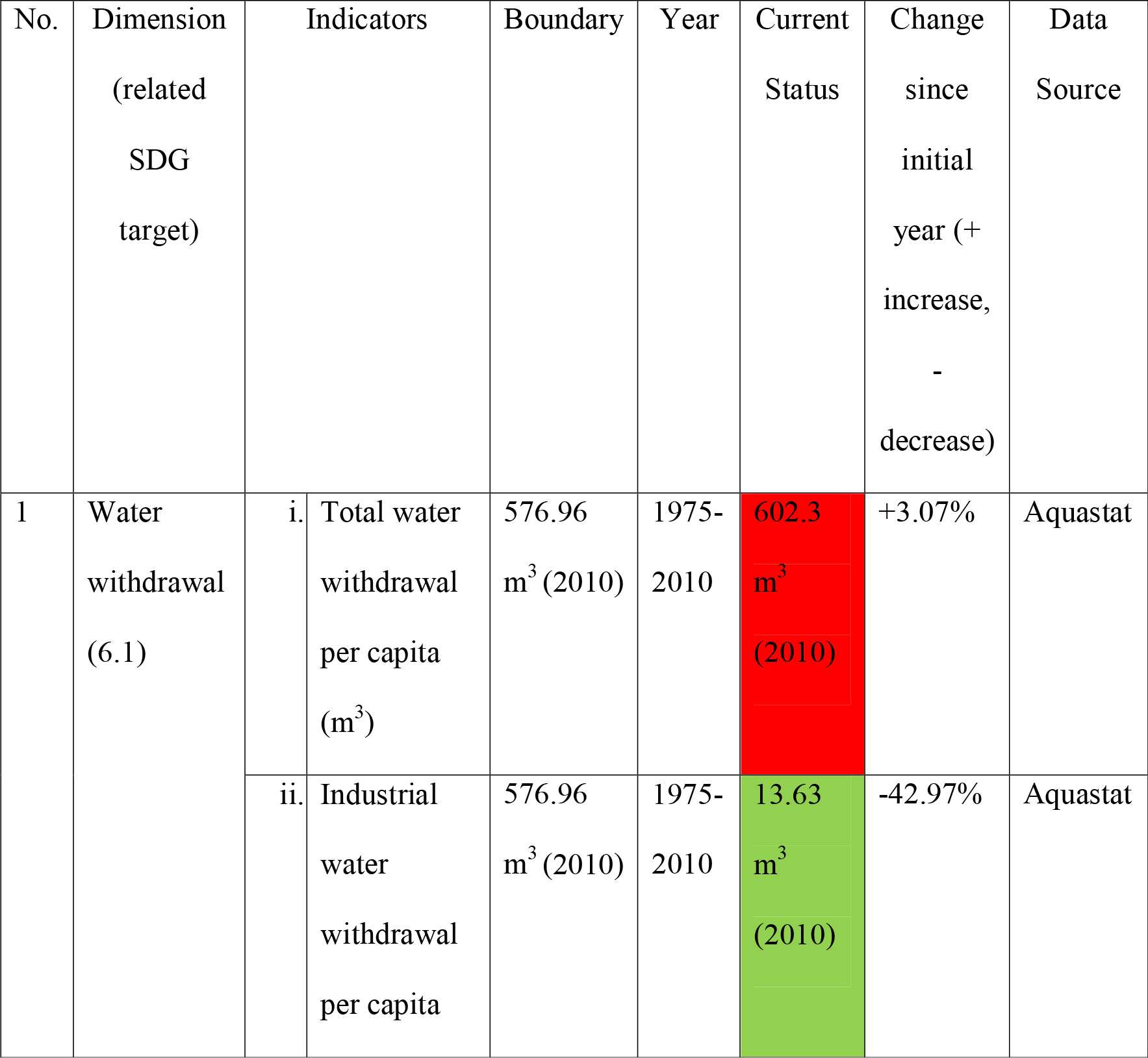
Biophysical indicators related to SDG 6 for India

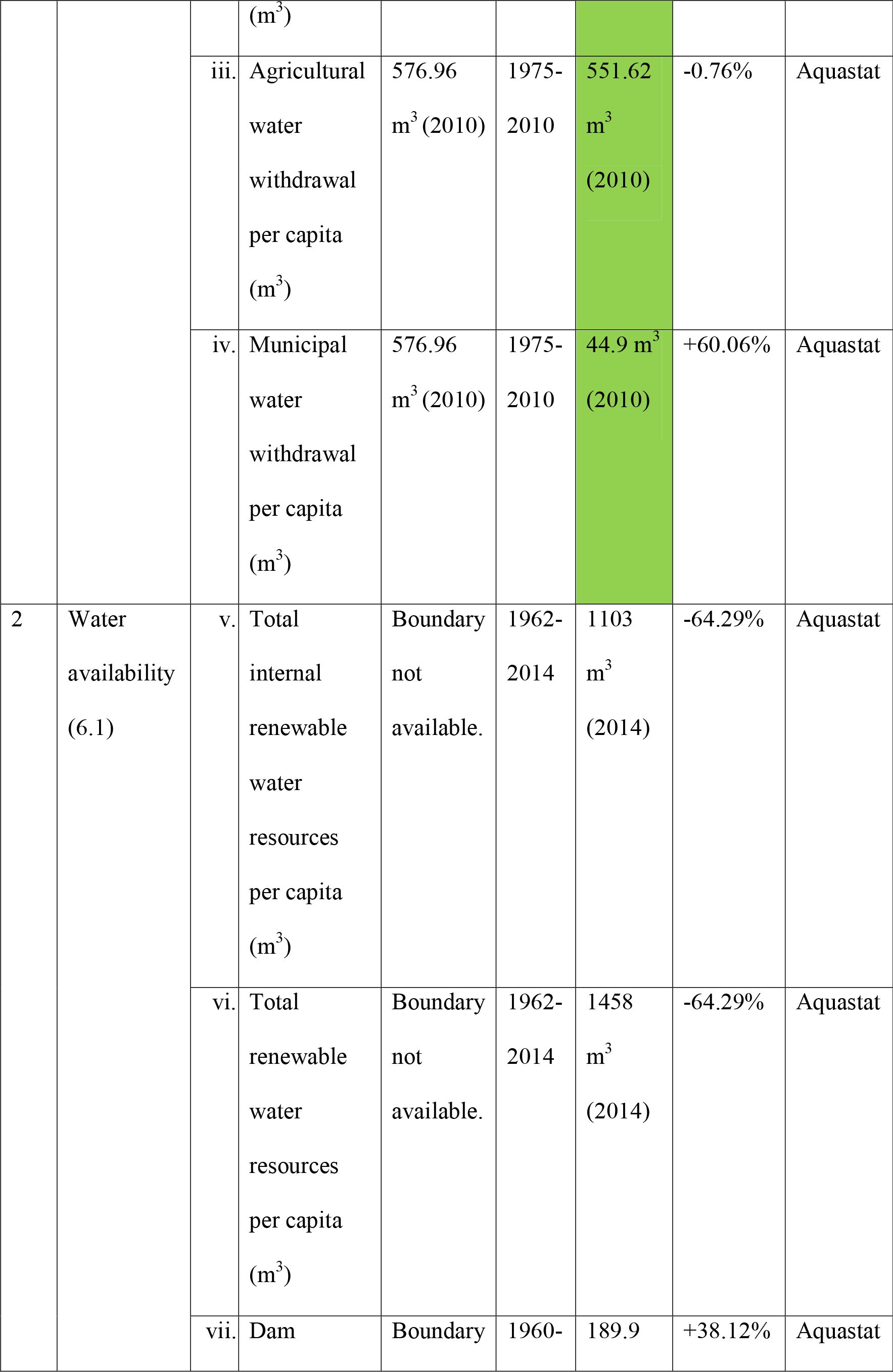

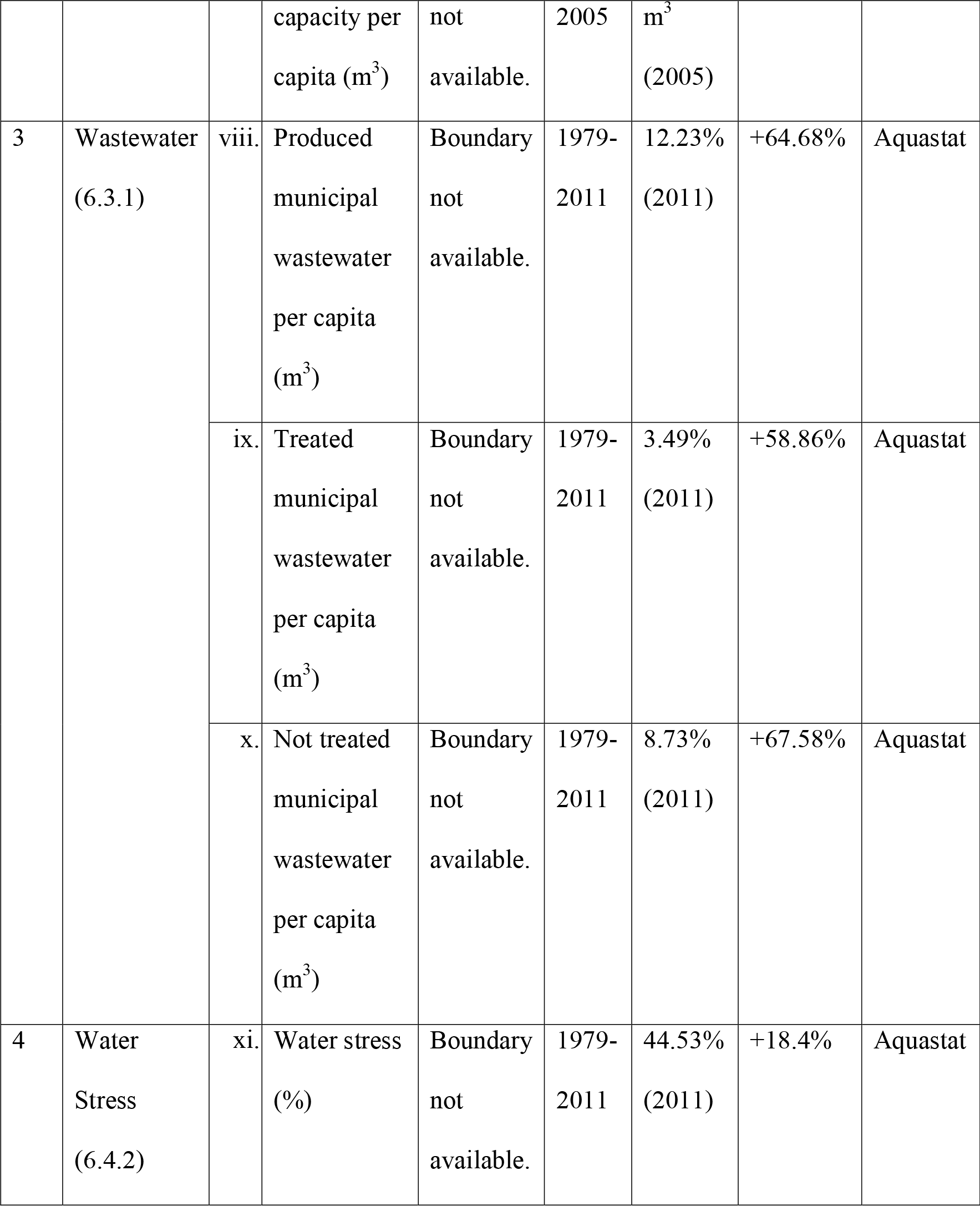

Values for indicators that are within per capita global average freshwater use PB are shown in green. Those have crossed the boundary, are shown in red.

### 2.2. Social development Indicators

We analysed 17 indicatorsunder7 dimensions that indicate social development aspects related to SDG 6 (Table 2). These 7 dimensions of social development indicators are - open defecation (3 indicators), basic drinking water (3 indicators), basic sanitation services (3 indicators), safely managed drinking water services (1 indicator), improved sanitation services (3 indicators), improved water source (3 indicators) and water use efficiency, WUE (1 indicator). We have also set a desirable threshold for social development indicators to get a better understanding of gaps between current status and desired status that can be reasonably accepted as the well-developed situation of SDG 6 for India. These are – less than 10% people for open defecation and 90% or more people for rest of 5 dimensions (excluding water use efficiency, WUE). We have also calculated the approximate time when desired thresholds were or will be met for social development indicators (excluding water use efficiency, WUE) using linear interpolation at business-as-usual, BAU scenario. The details of each indicator, along with duration, current status, change, threshold/boundary meeting time etc. are provided in Table 1 (biophysical indicators of SDG 6) and Table 2 (social development indicators of SDG 6).

**Table 2.**
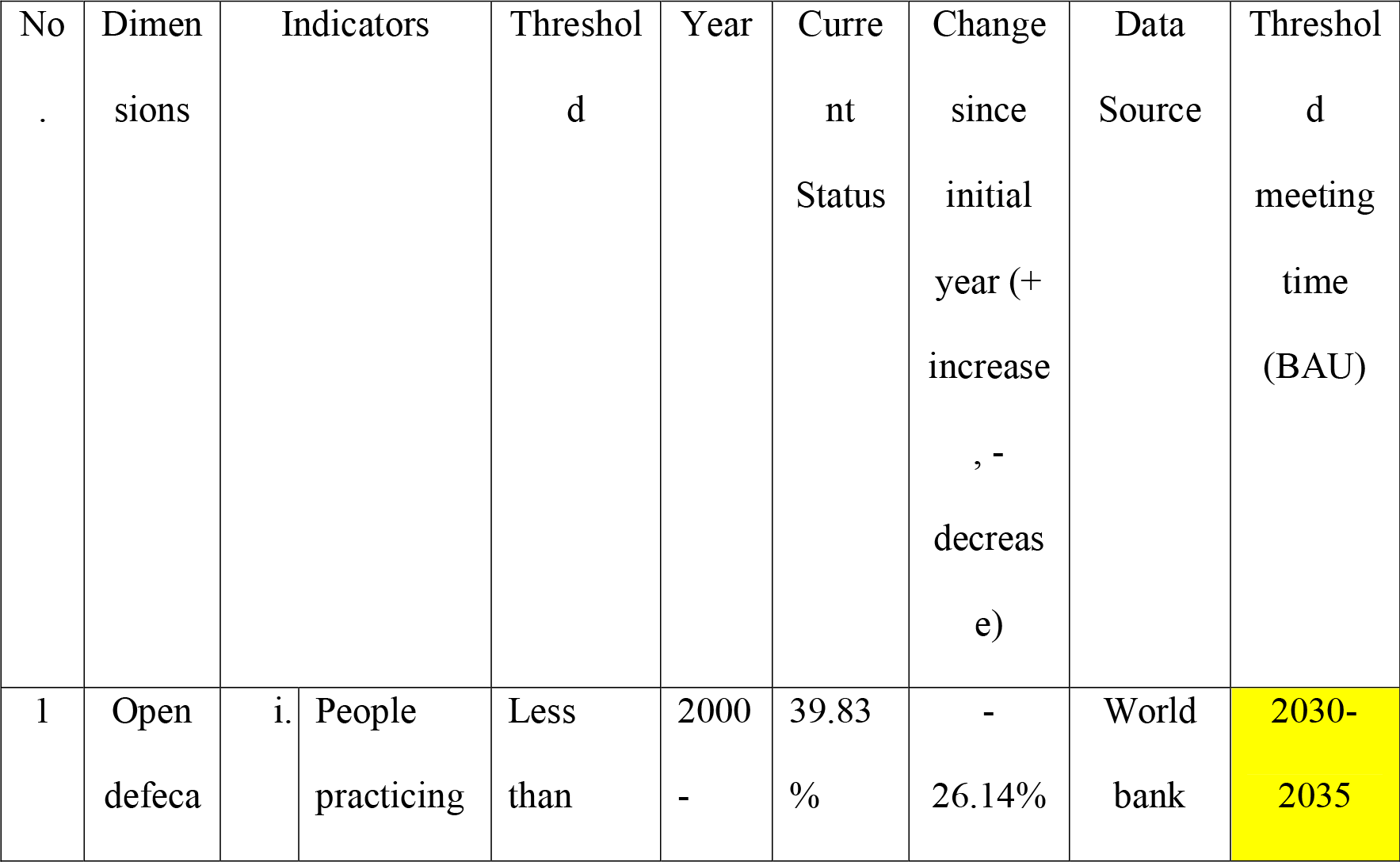
Social development indicators related to SDG 6 for India

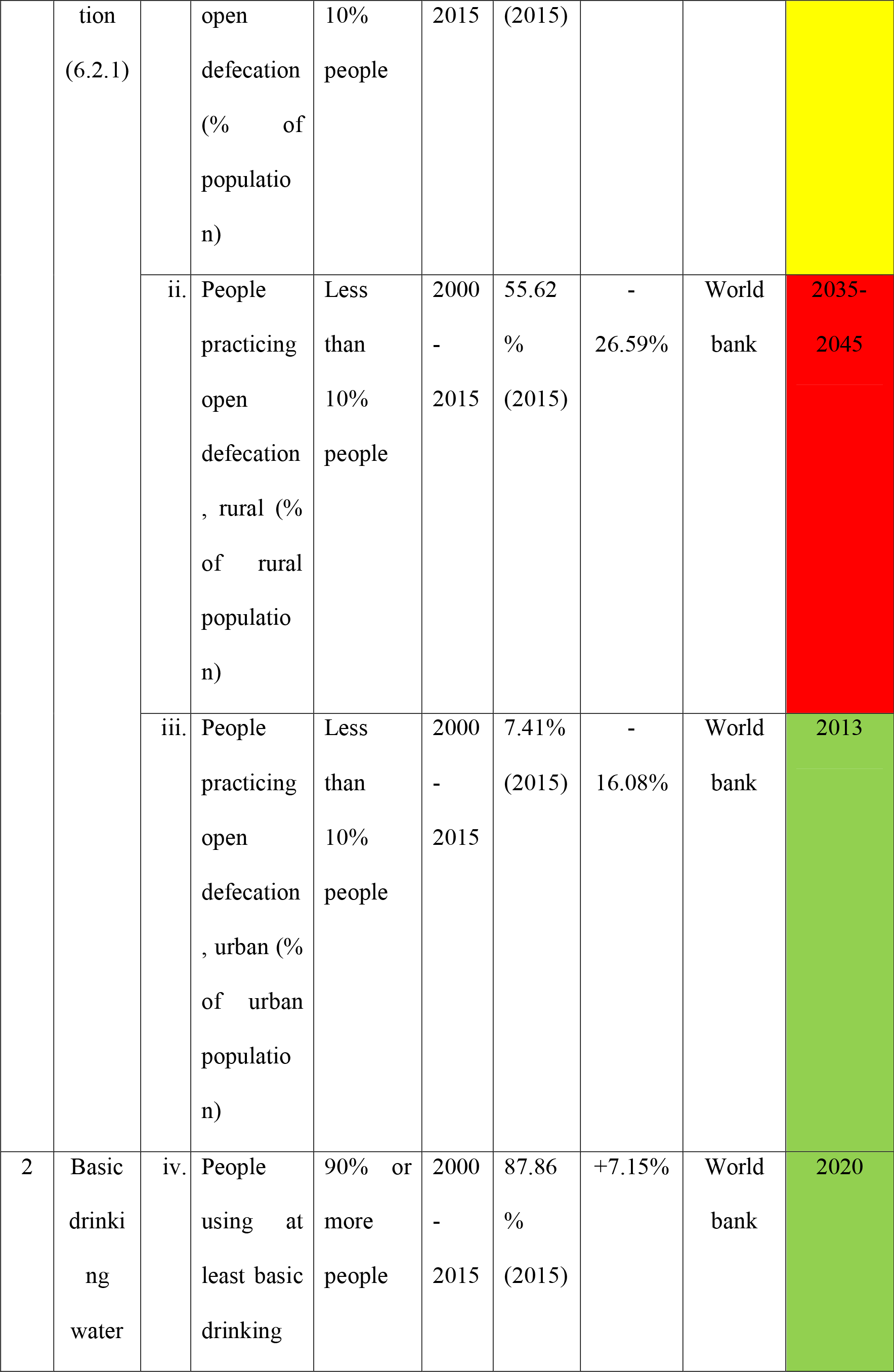

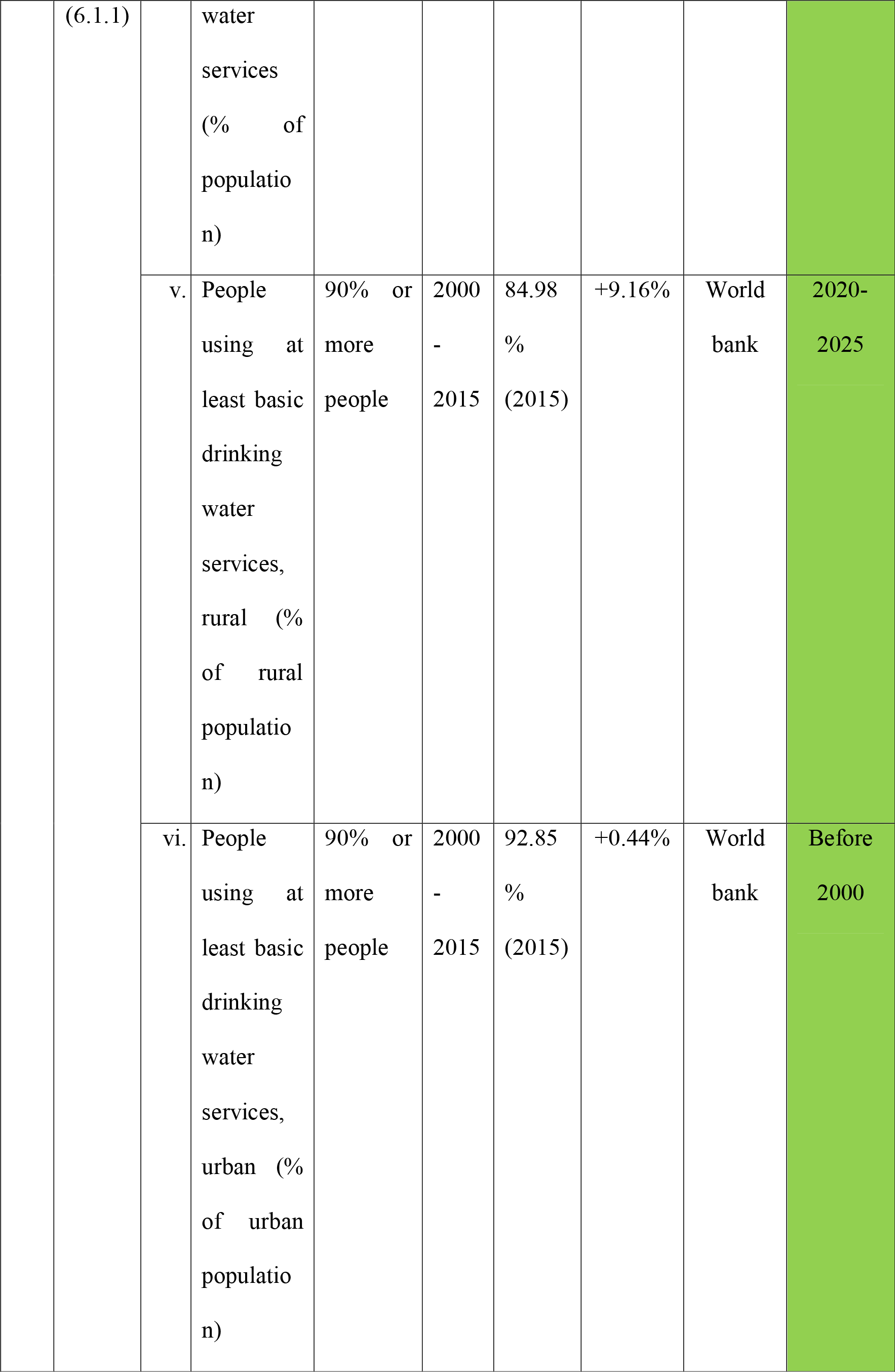

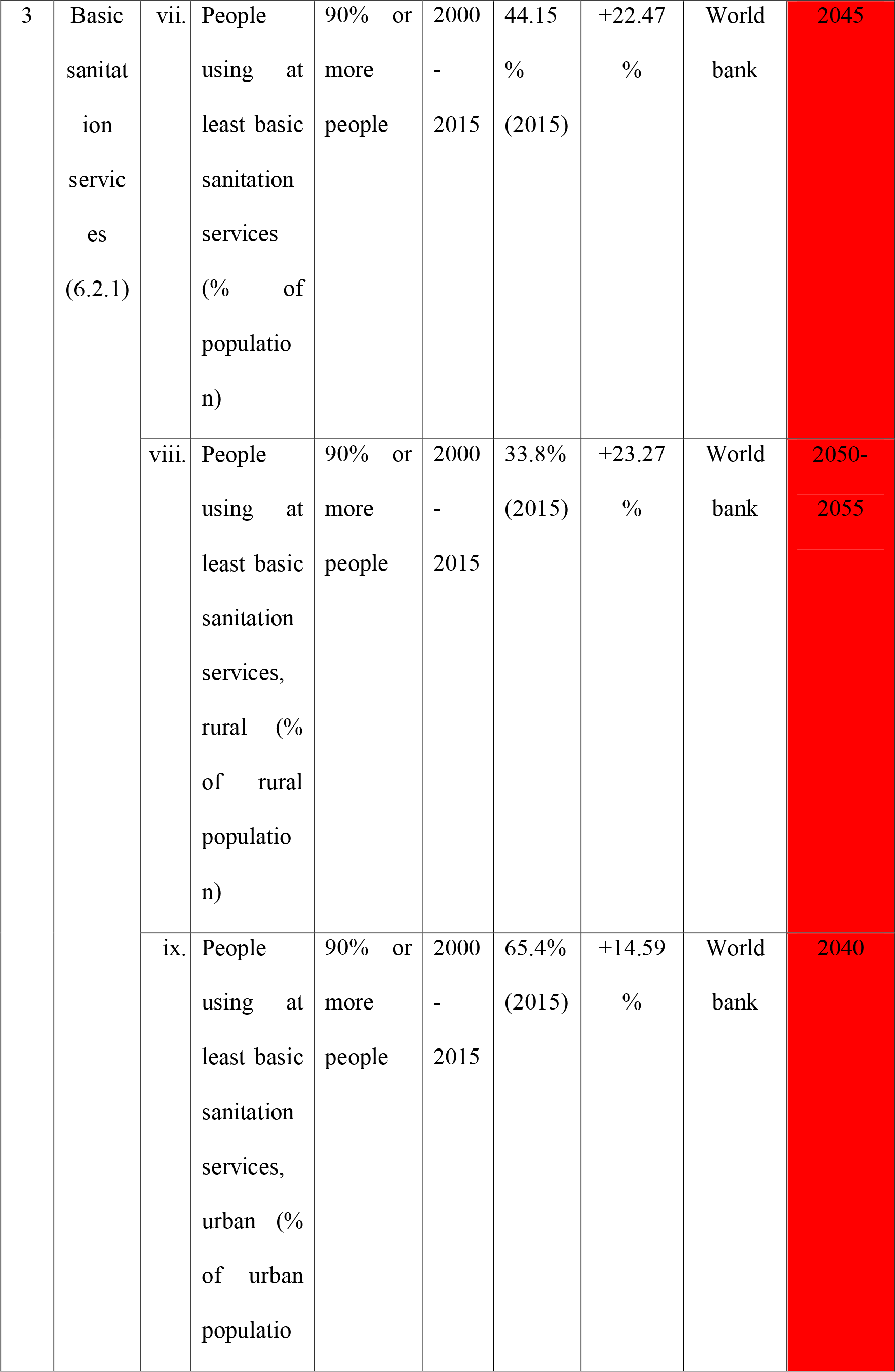

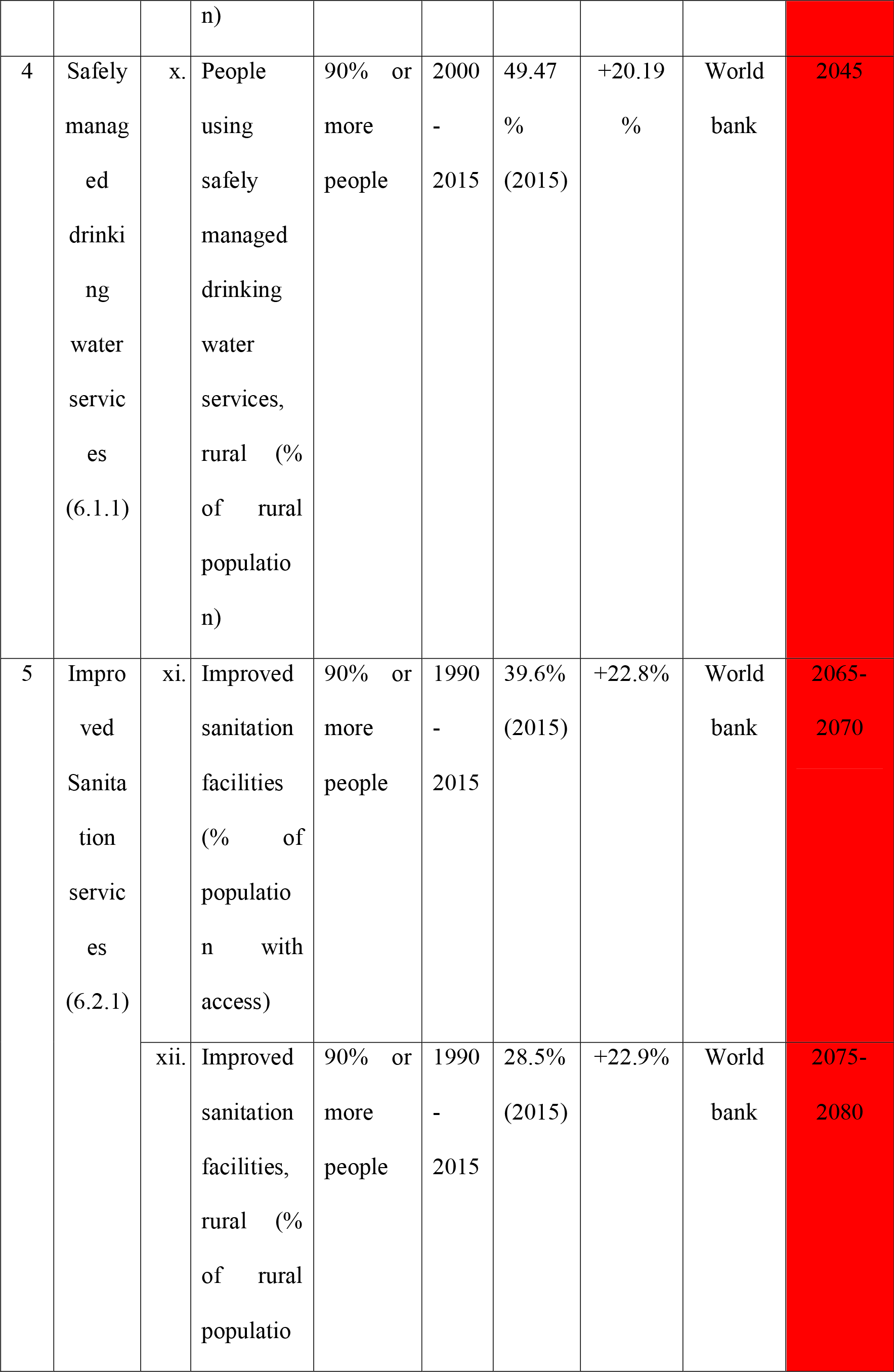

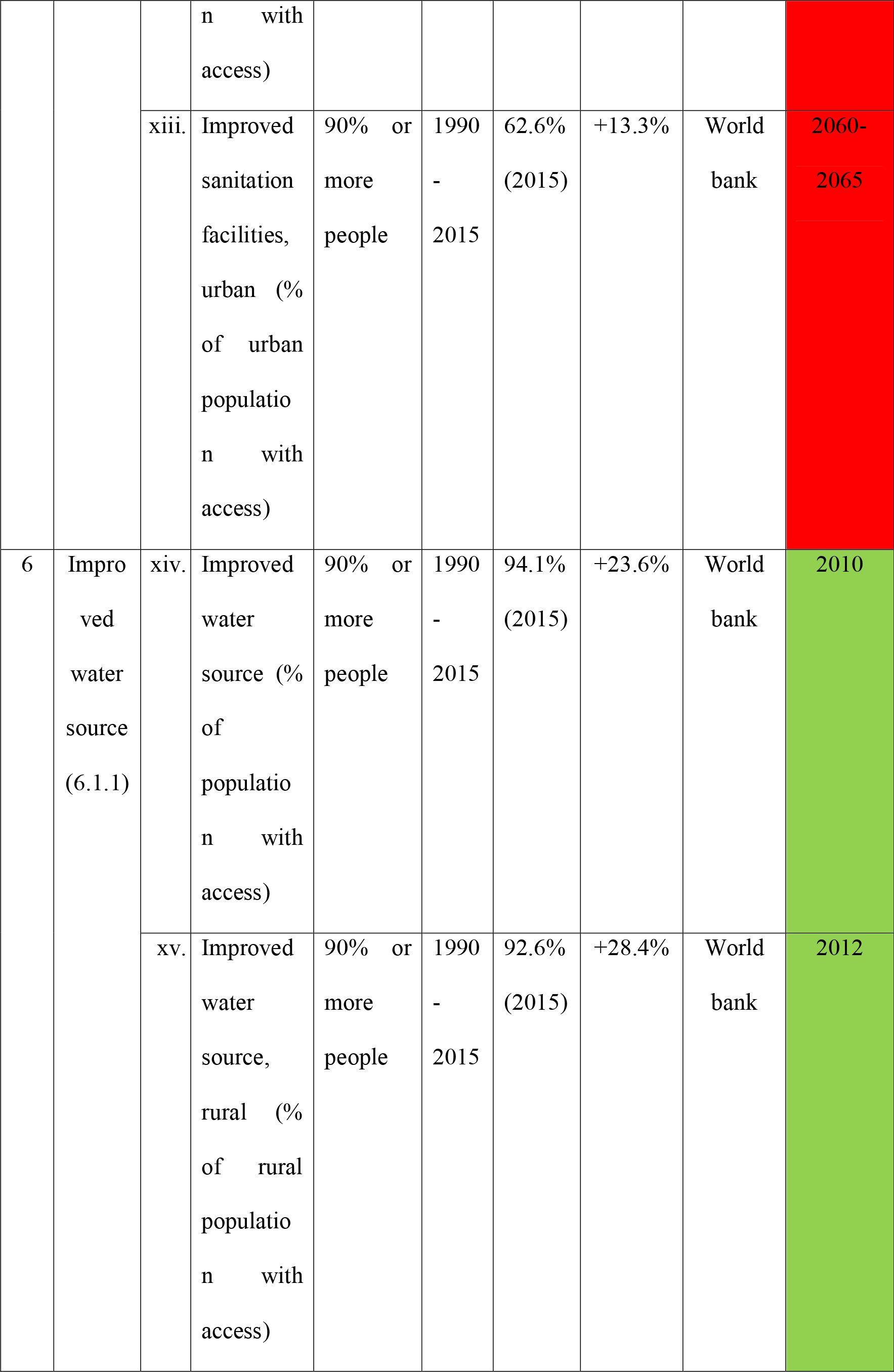

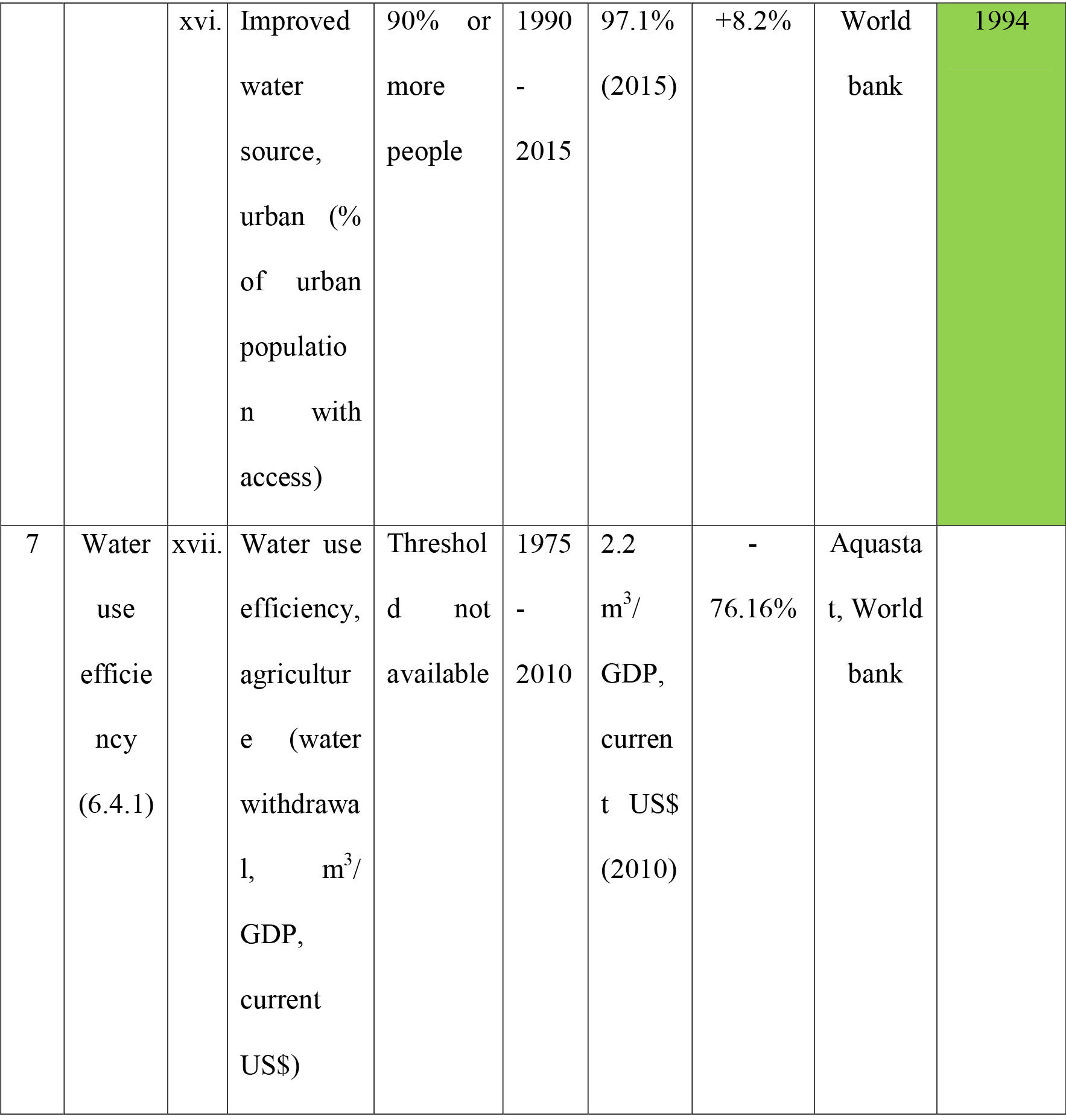

Indicators that are going to meet threshold within UN SGD target time (2030) are shown in green. Those going meet few years after that time are shown in yellow and indicators, which are going to meet desired threshold many years after 2030 are shown in red.

### 2.3. Planetary boundary of freshwater use

According to Rockström et al. (2009), the planetary boundary of freshwater use is the maximum withdrawal of 4000 km^3^ y^−1^ blue water from rivers, lakes, reservoirs, and renewable groundwater stores. We divided it with world population (from World Bank) to get per capita global scale boundary of 576.96km^3^ y^−1^ (2010).

### 2.4. Future scenario

As we have calculated most of the biophysical indicators on per capita basis (except – water stress, in %), it is possible to project probable future scenario of total consumption. We collected future population projection (2015-2050) data (median range prediction value of 50%) of India from UN DESA (2017 Revision) and then multiplied it with 10 per capita consumption indicators of SDG 6. We have calculated three projection series for each of 10 indicators of SDG 6, (i) with the lowest value that has happened in past year, (ii) highest value that has happened in past year and (iii) business-as-usual, BAU scenario with latest available data.

## 3. Results and Interpretation

### 3.1. Biophysical Indicators

Among the 11 indicators concerning biophysical aspects of SDG 6, 4 have decreased and 7 have increased over time (Table 1).

Both total internal renewable water resource and total renewable water resource have significantly decreased (64.29%) which is expected, as the population is growing but the available amount of renewable water resource is not. In future, with more population growth, per capita available renewable water resource is likely to decrease further. This problem can be tackled in 2 ways, (1) control of population growth in India and (2) increasing water productivity, i.e. perform more functions with less amount of water, in another word, becoming more water-efficient. Industrial water withdrawal has decreased significantly (42.97%) that might prove to be good, especially for India. More water will be available for agriculture (India has an agriculture-based economy) and domestic consumption (India is going to be one of the highest populated countries in the world). Per capita agricultural water withdrawal has decreased very less (0.76%) during the time (1975-2010), yet the population has increased much more i.e. total amount of agricultural water withdrawal have increased. It means water productivity in agriculture has not increased significantly in India. Therefore, increasing water productivity in agriculture i.e. growing more agricultural products using less water (direct or indirect use) should be one of the future concerns.

Total water withdrawal per capita have increased (3.07%). As population will increase in future, at this rate of increasing water withdrawal water scarcity will be more severe unless proper mitigation strategies are adopted. Municipal water withdrawal has also increased significantly (60.06%) which is probably due to increasing number of urban population. Thus, in 2050, if 50% of Indian population become urban dwelling (world urban prospects, UN-DESA, 2014 revision), this is likely to be continued, might even at higher rate. As a result, in future, it might not be possible to provide sufficient water for drinking and sanitation equitably to all urban people. Bringing down indiscriminate use of water in municipality areas is one of the probable solutions. If situation arise to more severe stage, water tax might be considered for implementation, especially for high-income urban localities. It is also a good time to start thinking about possible sources of freshwater in and around urban areas for supply to increased urban population in future. Continuous monitoring to check misuse of municipality provided water is needed. For domestic use in 2050, India would require 111 billion m^3^ water. Top 6 domestic water states (more than 8 billion m^3^ water) would be – Uttar Pradesh, Bihar, Maharashtra, West Bengal, Madhya Pradesh and Andhra Pradesh (IndiaStat 2018). (Supplementary Fig. 1). Total annual water requirement is increasing steadily in India. In 2050, water requirement is going to be almost three times the requirement of 1990. Water demand for power generation is going to increase more in 2025 and 2050 (IndiaStat 2018) (Supplementary Fig. 2). Dam capacity per capita have increased (38.12%). It means, as population is increasing, more numerous and larger capacity dams are being prepared or going to be needed to prepare in future to maintain this per capita rate. In one hand, more hydroelectric power is necessary for growing Indian population, in another hand, adverse effects of present and future construction of dams on aquatic life, especially riverine biota should be kept in mind (LeRoy Poff et al. 2007, Vörösmarty et al. 2010). Not only aquatic biodiversity is major source of livelihood of millions of people around river basins, but also protection and conservation of aquatic biodiversity is also priority. Produced and treated municipal wastewater has increased significantly (64.68% and 58.86%, respectively). But not treated municipal wastewater has increased more (67.58%). It means more amount of wastewater is remaining untreated than treated portion which brings bad effects of wastewater in environment, soil, biodiversity etc. along with it. More wastewater treatment plants or centres accompanied with more recycle and reuse of water are necessary in future. Water stress has also increased (18.4%). Current water stress level in India (44.53%) is almost 4 times more than average global level of water stress (12.82%). This indicates water stress is steadily increasing and would become more complex problem in future with growing number of Indian population. Changes of biophysical consumption indicators are depicted in Fig 2.

**Fig. 2.**
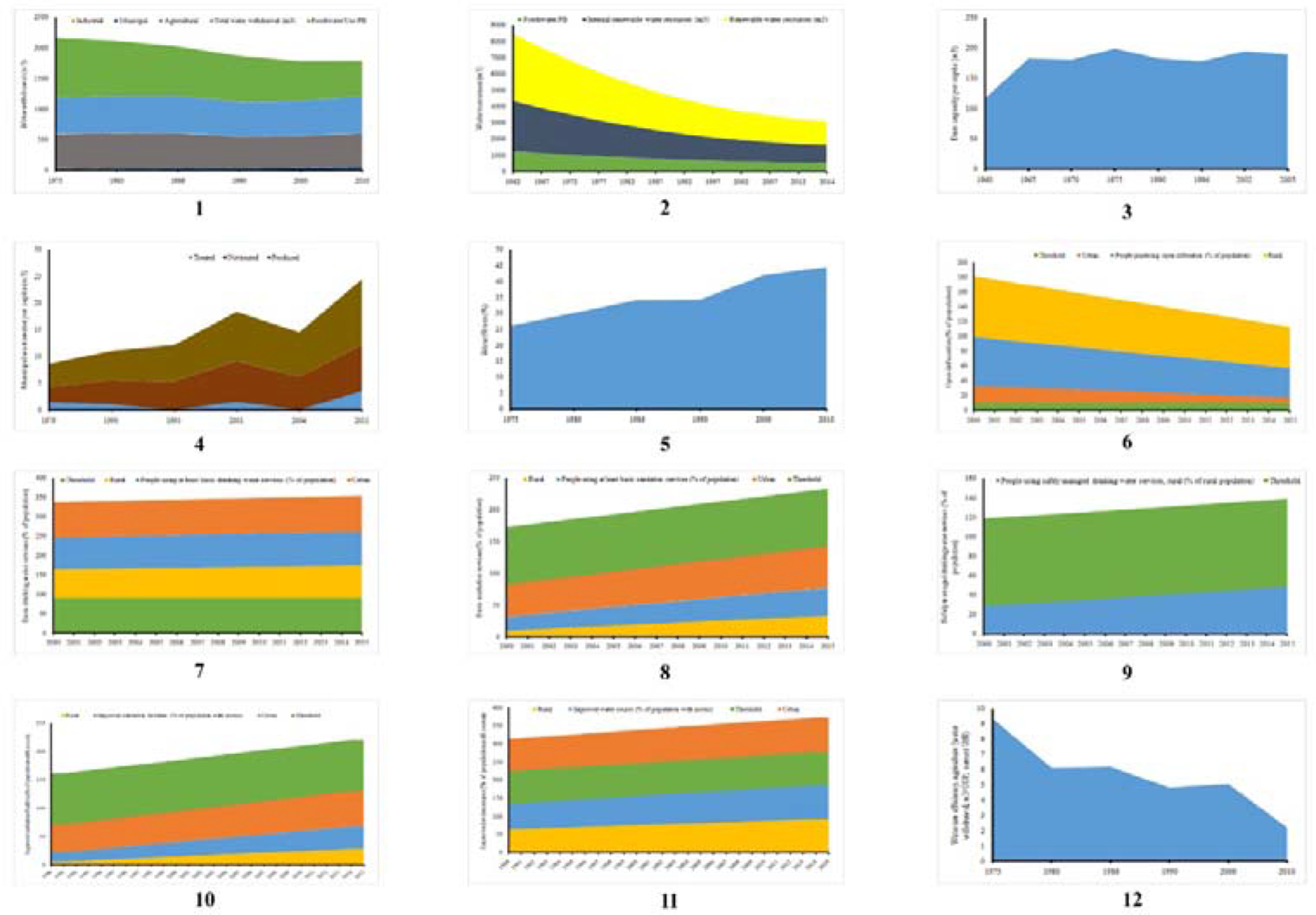
Changes in per capita biophysical indicators (1-5) and indicators of social development (6-12) related to sustainable development goal 6 in India with time. Green indicates global per capita boundaries for biophysical indicators and thresholds for indicators of social development. Biophysical indicators are - (1) water withdrawal (m^3^), (2) water resources (m^3^), (3) dam capacity (m^3^), (4) municipal wastewater ((m^3^) and (5) water stress (%). Indicators for social development are - (6) open defecation (%), (7) basic drinking water services (%), (8) basic sanitation services (%), (9) safely managed drinking water services (%), (10) improved sanitation facilities (%) (11) improved water source (%) and (12) water use efficiency in agriculture (m^3^/GDP, current US$).

### 3.2. Social development Indicators

Of the 17 indicators of social development aspects of SDG 6, 4 have decreased and 13 have increased over time (Table 2).

Open defecation rate in India has decreased in all 3 cases, overall (26.14%), rural (26.59%) and urban (16.08%). Urban open defecation rate has already met with the desired threshold (10% or less) in 2013. Although the success, it should be also kept in mind that with increasing number of urban populous in coming decades, maintaining this rate might prove to be tougher. So, necessary actions should be made parallelly for implementation in need be in future. Remaining two will not be able to meet SDG criteria at this rate within 2030 (2030-35 and 2035-40, respectively). It means open defecation in rural areas needs to decrease more. There are 2 possible ways for that, (1) more public and household toilets are to be constructed in rural areas and (2) spreading awareness about adverse effects of open defecation on environment and health in rural areas. On average, 35.09% villages in India are open-defecation free as of 2017. Four are lowest (0%) among them (Dadra and Nagar Haveli, Daman and Diu, Goa and Puducherry) and five have highest (100%) achievement (Haryana, Himachal Pradesh, Kerala, Sikkim and Uttarakhand) (Supplementary Fig. 3) (IndiaStat 2018). Water use efficiency in agriculture has also decreased significantly (76.16%) which is good, i.e. less amount of water is being withdrawn for the same amount of GDP (in current US$, 2010).

Basic drinking water services has also increased (overall, 7.15%; rural, 9.16% and urban, 0.44%). Basic drinking water in urban areas has already met with a desirable threshold (90% or more) before 2000. Remaining two will also be able to meet SDG criteria at this rate within 2030 (2020 and 2020-25, respectively). Basic sanitation services have increased in India (overall, 22.47%; rural, 23.27% and urban, 14.59%). But, none of the 3 will be able to meet desired SDG criteria within 2030 (2045, 2050-2055 and 2040 respectively). Immediate steps are needed to improve awareness and basic sanitation system. Safely managed drinking water services in rural Indian population has also increased over time (20.19%). But it will also not be able to meet desired SDG threshold within 2030 (2045). Improved sanitation facilities have also increased over time (overall, 22.8%; rural, 22.9% and urban, 13.3%). As none of the 3 indicators will be able to meet desired SDG criteria within 2030 (2065-2070, 2075-2080 and 2060-2065 respectively), improvement of sanitation services requires serious attention in India (both rural and urban area). Improved water source has increased over time (overall, 23.6%; rural, 28.4% and urban, 8.2%). All of these 3 indicators have already meet desired SDG criteria (2010, 2012 and 1994 respectively). To sum up, in India, among 16 indicators with desirable thresholds, 7 have either already met or going to meet SDG criteria within 2030, 1 indicator will reach the target just after 2030, and remaining 8, at current rate, will be able to meet in distant future (2040-2080), requiring immediate attention. Changes in social development indicators over time are seen in Fig 2.

### 3.3. Planetary boundary of freshwater use at the national scale

From this analysis, we can see that Indian per capita total water withdrawal (2010) has already crossed global average freshwater use PB (576.96 m^3^, 2010) (Fig. 1). This means that with growing population it would be tougher for to achieve UN SDG 6 targets remaining under safe biophysical limits of water resource use.

### 3.4. Future of SDG 6

Fig 3 shows total amount of probable biophysical consumption up to 2050 in 9 graphs. In 2050, if lowest per capita rate can be maintained, total water withdrawal would increase 18.98%, at a business-as-usual rate, it would increase 25.8% and at highest per capita rate, it would increase 26.57%. It means even with grown population level of 2050, it is possible to lower the rate of total water withdrawal 6.8-7.5% and which is a significant amount. In agricultural water withdrawal in 2050, at lowest per capita rate, it would increase 20.98%, at a business-as-usual rate, it would increase 25.8% and at highest per capita rate, it would increase 29.1%. Agricultural water withdrawal might to be 1.4 times more in 2050 than 2010. For industrial water withdrawal, in 2050, at lowest per capita rate, it would decrease 7.72% which is a very positive scenario. But at a business-as-usual rate, industrial water withdrawal would increase 25.8% and at highest per capita rate, it would increase 57.7% in 2050. For municipal water withdrawal, at lowest per capita rate, it would decrease 46.18% in 2050 which is a positive scenario, whereas for business-as-usual (this is also highest per capita) rate, it would increase 25.8%. For produced municipal wastewater, in 2050, it is possible able to reduce it to 53.02% maintaining lowest per capita rate. Otherwise, at business-as-usual (highest per capita) rate, it would increase 24.82%. Municipal wastewater production might be 1.3 times more in 2050 than 2011. For treated municipal wastewater, it would decrease to 60.71% maintaining lowest per capita rate. But at business-as-usual (highest per capita) rate of wastewater treatment, amount of treated municipal wastewater would increase 24.82% which is also positive scenario. For non-treated municipal wastewater, it is possible to decrease it to 56.83% maintaining lowest per capita rate in 2050. On another hand, at business-as-usual (highest per capita) rate, it would increase 24.82% which paints a bleak future of India. This non-treated municipal wastewater might be 1.33 times more in 2050 than 2011. Both total renewable water resource and total internal renewable water resource, if lowest per capita (which is also business-as-usual) rate can be maintained by India, only 22% more renewable water resource will be needed in 2050. Otherwise, it would plunge to 72.15% if highest rate of renewable water resource per capita is to be maintained.

**Fig. 3.**
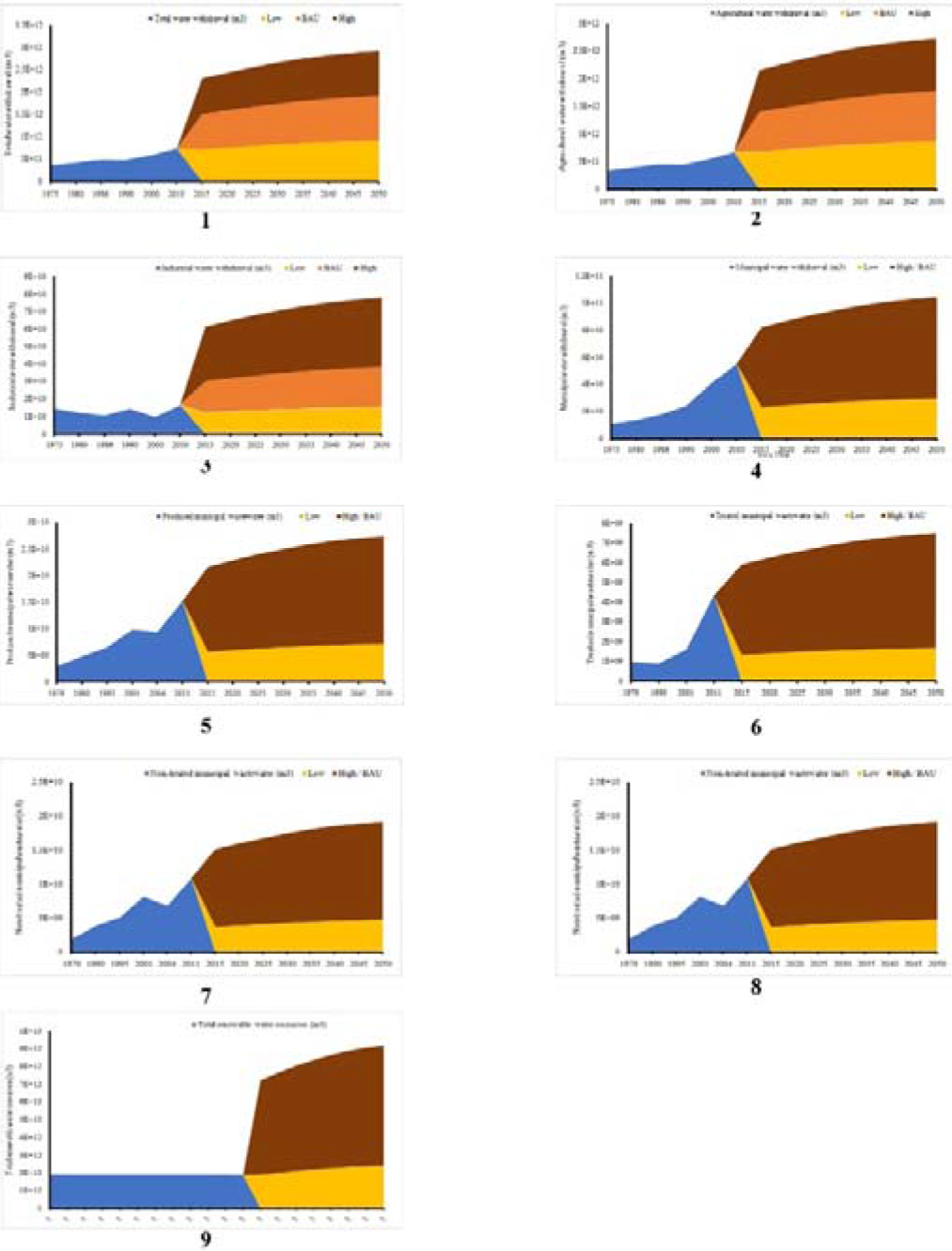
Future scenario of indicators of biophysical consumption related to sustainable development goal 6 for India up to 2050. Blue indicates changes in total values of consumption. Brown indicates projected total values at the highest rate of per capita consumption; Orange indicates projected total values at business-as-usual (BAU) rate of per capita consumption, Yellow indicates projected total values at a lowest rate of per capita consumption.

## 4. Discussion

This study clearly shows that the per capita biophysical resources are decreasing over time. This trend would continue further with respect to population explosion and growing urbanisation trends in India. However, it also shows that social developments related to water and sanitation are improving, clearly at the cost of environment. To provide positive social development and maintaining the same while the population is exploding will be tough to handle for the government. Especially, if all of these developments come at the cost of further degradation of the environment.

During this study, we confronted some problems along with their probable mitigation strategies. First, availability of biophysical resource consumption data is not good enough. For the same reason, we were unable to perform any statistical analysis. In future, analysis and freely available data concerning biophysical consumption at state and district level should be prepared and maintained, both for general awareness and scientific research purposes. Second, long-term data are not available for most of the indicators which are necessary for more accurate projection of future trends. Third, neither state-level comprehensive data for most of the indicators are available, nor the district or any other lower unit level. This simply creates a homogeneous outline of resource consumption and access at national level. This is not enough to redesign policy framework or restructure governance in accordance with scientific findings. Though it is enough for preliminary studies, would not be enough in future, especially when adaptive policy needs to be implemented customized for the local context. Fourth, abundance and availability of data of different biophysical parameters are much more than different socioeconomic indicators. It shows a clear bias towards the maintenance of biophysical data. This is not suitable for contemporary research works and creates problem to assess sustainability for all three dimensions (i.e. environment, society and development). Fifth, the set of indicators for sustainable development of water and sanitation sector does not yield a very complete and comprehensive picture of SDG 6 in India, especially in view of the socioeconomic factors that exist within the country and the regional biophysical resource consumption that interact with them. Therefore, there is ample scope for adapting the indicators to better suit India’s specific needs in terms of reflecting both regional and national scenario. To this end, additional indicators could be developed.

Sixth, as this study only etches the surface of multi-indicator multilevel continuous monitoring of SDG 6, it is not rigorous enough to map synergy and trade-off between SDG 6 and other sustainable development goals (Griggs et al. 2014, Nilsson et al. 2016). Seventh, awareness and know-how about water resource management in India is only catalysed by the government in broad scale. Thus, government, nongovernment organizations (NGOs) and the public, together should take more proactive steps towards promotion and utilization of water resources to achieve sustainable development goals in water and sanitation (i.e. SDG 6) in India. Eighth, to control misutilization and overexploitation of water resources, especially in urban areas, regulatory instruments, such as - water utilization taxes, water recycling taxes etc. will remain a vital component of policies, as efforts to increase water efficiency and reduce environmental and socioeconomic impacts are usually unsupportable against a background of low cost availability. Ninth, if consumers do not purchase devices with improved water use efficiency and environment characteristics (e.g. agricultural transport and household appliances etc.), when such devices are available commercially, these products should be supplemented by more direct government intervention and investment if advancement toward sustainable development in water resources are to be made. Tenth, since agriculture and household are two major water consuming sector in India, utilization of recycled and treated wastewater should be advocated. Eleventh, since agricultural water withdrawal is highest in India among all other sectors, increasing agricultural water use efficiency should be the highest priority through drip irrigation, hydroponics, urban terrace and rooftop agriculture, utilization of treated household wastewater in agriculture etc.

Water resources and its utilization are intimately connected to sustainable development. For Indian societies to attain sustainable development in water and sanitation, much effort must be devoted not only to discover sustainable ways to consume of water resources but also to increase the water use efficiencies (WUE) of processes utilizing water resources. Also, correlated environmental concerns must be addressed, continuously monitored and any type or degree of declination must be solved. This proposed research framework would progress our current understanding and measurement of SDG 6. Through measurement and identification of how, when, and why changes in biophysical aspects of water and sanitation influence human socioeconomic conditions over both time and space and vice-versa, we can more fully appropriate and anticipate the impact of the bidirectional nature of human-water interactions. This understanding might also be used to monitor, update and refine the national SDG 6 framework. This would arm natural resource planning authorities with a decision support and policy management tool that more accurately reflects the trade-offs between various components of the SDG 6. As a final note, to make India water secure and sustainable in 2030, India must endeavour to increase its efforts to attain greater efficiency in utilization, consumption of water for both biophysical and socioeconomic purposes.

## Acknowledgement

This study was supported by FRPDF scheme (2013-2017) of Presidency University, Kolkata. We would like to thank Sk. Rohan Tanvir, The Institution of Engineers (India) for his kind assistance during the preparation of diagrams.

